# Compression for population genetic data through finite-state entropy

**DOI:** 10.1101/2021.02.17.431713

**Authors:** Winfield Chen, Lloyd T. Elliott

## Abstract

We improve the efficiency of population genetic file formats and GWAS computation by leveraging the distribution of sample ordering in population-level genetic data. We identify conditional exchangeability of these data, recommending finite state entropy algorithms as an arithmetic code naturally suited to population genetic data. We show between 10% and 40% speed and size improvements over dictionary compression methods for population genetic data such as *Zstd* and *Zlib* in computation and and decompression tasks. We provide a prototype for genome-wide association study with finite state entropy compression demonstrating significant space saving and speed comparable to the state-of-the-art.

## 1 Introduction

A full description of the mapping from genotype to phenotype in humans will be a centerpiece of science, and will lead to an understanding of the human condition, and will lead to advances in medicine. Genome-wide association study (GWAS) is currently the main tool used to create, and revise *draft* versions of this genotype-to-phenotype mapping. The pursuit of such studies is a world-wide effort conducted and considered by labs in almost every university and also in biotechnology and pharmaceutical companies (Visscher et al., 2017). The computational resources required for GWAS are large, and so improving the efficiency of the manipulation and storage of genetic data and of conducting GWAS will increase the pace and accessibility of GWAS research, and also allow more efficient use of public and private funding.

To this end, researchers have made improvements to the compression of genetic data, including support for the cutting-edge *Zlib* (Gailly and Adler, 2004) and *Zstd* (Collet and Skibinski, 2019) libraries in the popular *bgen* and *ped* formats (Band and Marchini, 2018; Chang et al., 2015). This has allowed cheaper dissemination and storage of large-scale consortia such as the UK Biobank (Band and Marchini, 2018; Bycroft et al., 2018). The *bgen* and the new *pgen* formats have also been extended to provide variable or low bitrates for lossy compression of dosages for large genetic datasets (Band and Marchini, 2018; Chang *et al*., 2019).

In this work, we further improve the compression of population-level genetic data using ideas stemming from recent advances in source coding (Shannon, 1948) including finite state entropy (Collet, 2019) and asymmetric codes (Duda, 2013). These recent advances are arithmetic codes (Huffman, 1952) for compressing streams of conditionally independent symbols and show improved performance in size and speed of population genetic data compression tasks.

### 1.1 Conditional independence in genetics

File formats for population level genetic data such as *bgen* and *plink2*’s *ped* and *pgen* formats store genotypes in compressed variant blocks (Chang et al., 2015), in which a bitstring representation of the genotypes for all subjects are compressed separately for each marker (*i.e*., they are stored in a variant-major form). This organisation is optimal for the purpose of fast GWAS. However, the samples collected in population-level genetic data are usually exchangeable, or partially exchangeable (Orbanz and Teh, 2010). This exchangeability arises because genotypes for subjects are stored in a fixed order, and this order is identical for all variants. When a study is conducted on a homogeneous population, this fixed order does not provide information about the genotypes, leading to conditional independence among the genotypes at a fixed variant (*i.e*., exchangeability). The independence is conditioned on the minor allele frequency (*a.k.a*. the MAF), or other variant specific measures. If a pedigree, or a genetic similarity matrix (Patterson et al., 2006), or population indicators is also considered, then the genotypes are jointly exchangeable (*i.e*., the joint distribution on the genotypes is still exchangeable and invariant under permutation, as long as the permutation is also applied to the pedigree, the genetic similarity matrix, or the population indicators).

For stratified or related populations, often the fixed order still provides no information about the genotypes (the subjects may be presented in an order given by sorted and random subject identification numbers) But even if a heterogeneous sample is considered in which the order indicates subpopulation identity in a block structure (leading to partial exchangeability), conditional independence still exists among all subjects in each subpopulation, and the number of blocks in the structure is often small compared to the number of samples. These considerations imply that a bitstring representation of the ordered genotypes is well approximated by independent and identically distributed draws from a fixed law:

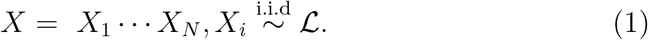

Here *X* is the bitstring for a marker typed for *N* subjects and *X*_i_ is the bitstring for subject *i* and 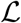 is a discrete law on floating point numbers or vectors of floating point numbers (the floating points are relevant for variants that include dosage information along with discrete genotypes). For example, if *X* involves hard-called genotypes with MAF *p* under the Hardy-Weinberg equilibrium (*a.k.a*. HWE), then *X*_i_ is a bitstring representation of the set {0, 1, 2} and 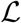 is the law described by a discrete distribution taking the value 0 with probability (1 — *p*)^2^ and 1 with probability 2*p*(1 — *p*) and 2 with probability *p*^2^ (here the level of *X*_i_ is the number of minor alleles). In other situations, such as for data in which dosages or genotype likelihoods or phase are recorded, 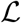 and the support of *X*_i_ can be suitably modified to still present data in the form (1).

Population-level genetic file formats often use *Zstd*, *Zlib* or *gzip* or similar ‘dictionary’ style compression methods for compression of their variant blocks. These methods work by building a dictionary of strings that occur commonly in the uncompressed bitstring and then replacing those aspects of the bitstring with keys into a dictionary (Sayood, 2012). However, due to the exchangeability or partial exchangeability of bitstrings of the form (1), for a dictionary value to be *viable*, many of the permutations of the dictionary value around the boundaries of the bitstring representation of a single genotype must appear in the dataset. We call this the ‘exploding dictionary’ problem. Here, when we say a dictionary value is *viable* we loosely mean that the value occurs often enough in the uncompressed data that its inclusion in the dictionary decreases the size of the compressed bitstring. Roughly speaking, the independence displayed in (1) reduces the extent to which dictionary values are viable. On the other hand, source coding theory (Shannon, 1948) is designed to describe and compress bitstrings that display independence such as (1). Our method, which we refer to as *agent* (arithmetic codes for *genet*ic data), exploits this exchangeability and independence by specifying the compression for genotype blocks with such source coding theory.

### 1.2 Preliminary demonstration

In Figure 1 we consider simulation replicates involving half a million samples and 1 variant, with MAF varying from 0.01 to 0.5, and we assume HWE and a random subject ordering (*i.e*., full exchangeability). We use reference implementations for both *fse* and *Zstd* (Collet, 2019; Collet and Skibinski, 2019). We show improved performance in both space and runtime for the arithmetic code at all MAF levels. The largest runtime improvements are seen at high MAF (Figure 1 *Top*) and the largest size improvements are seen at low MAF (Figure 1 *Bottom*). Notably, we find an apparently constant relationship between runtime and MAF in the *fse* arithmetic code. Theoretically, the runtime must be increasing with the size of the compressed variant block (even as it appears constant), as this block must be written to RAM (random access memory) or disc, and also the size of the finite state machine used by *fse* must also increase with MAF. But however, Figure 1 indicates that these increases may be amortized. The *fse* method improves upon *Zstd* in runtime by 8% to 34% and in size by 52% to 26%, as the MAF is varied.

**Figure 1:**
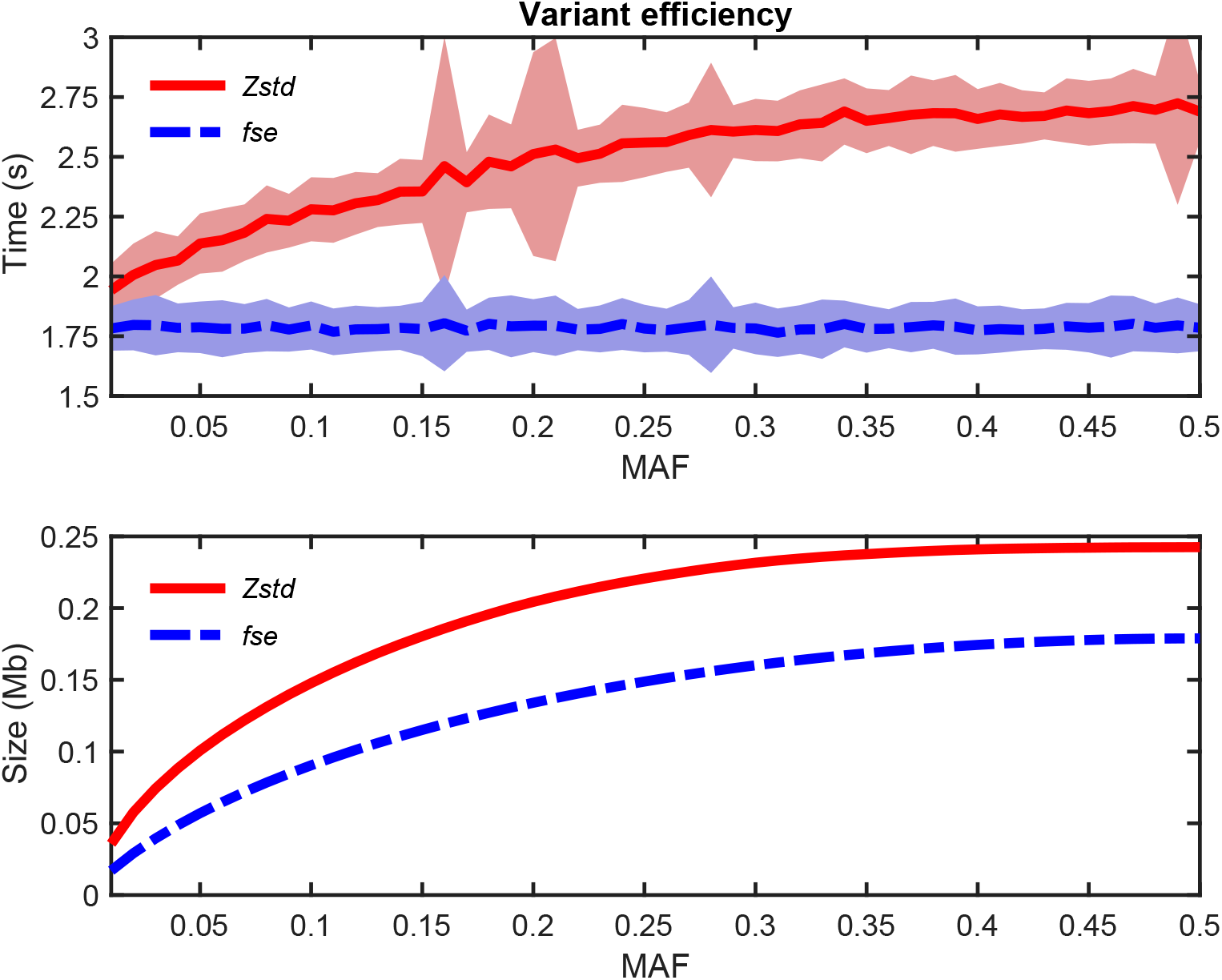
Computational efficiency of encoding a single variant simulated for one million subjects with a given minor allele frequency (MAF) using *Zstd* or *finite state entropy* (*fse*), with 120 replicates per condition. *Top*) Runtime of encoding in seconds (y-axis) and one standard deviation. *Bottom*) Size of encoding in megabytes (y-axis). Improved performance of *fse* over *Zstd* is demonstrated.

### 1.3 Related work

The variant-major organisation of file formats in population-level genetics allows random access to variants, which is required for optimised GWAS and PHEWAS (phenotype-wide association study). In another line of recent work, researchers have used linkage between adjacent variants to decrease compression size using trees, and Burrows-Wheeler transformations (Adjeroh et al., 2002; Kelleher et al., 2018). We do not consider a comparison with these works, as they do not allow random access.

We also note that recent advances in compressing genomes (for example the Nucleotide Archival Format Kryukov et al.; 2019) and alignment-free genomes Sweeten (2019) have lead to excellent compressive bitrates for use in improved storage of *de novo* genomes and computation of distances between sequences, with application to compressing assembled genomes. Recent advances in compressing assembled genomes also include Liu et al. (2017), which also involves arithmetic codes. The *fse* algorithm has also been used in this setting (Holmes, 2016). However, these works are not relevant for the context-free nature of the variants (such as single-nucleotide polymorphisms, or SNPs) considered in population-level genetic data. To our knowledge this manuscript is the first work to highlight the exchangeable or partially exchangeable nature of population genetic data and recognise that algorithmic codes (including *fse*) are most appropriate for compression of variant-major stores.

In Section 2, we review the *fse* and *ans* methods (summarizing Duda 2013 and Collet 2019). In Section 3, we provide runtime and filesize results for the *agent* tool on the Thousand Genomes Project (The 1000 Genomes Project Consortium, 2015) and simulated genomes.

## 2 Approach

*Finite state entropy* (*fse*) is a member of a family of general asymmetric numerical systems (*ans*) known as *tabled asymmetric numeral systems* (*tans*), described in Duda 2013. These methods combine the advantageous compression ratio of arithmetic coding with the favourable speed of Huffman coding. To fully understand the justification and advantages of *fse* and *tans* methods in general, we briefly summarize the developments leading to the state-of-the-art performance of these methods.

### 2.1 Tree-based coding and the integer bit problem

The original compression algorithm described in Shannon 1948 (Shannon-Fano coding) represents symbols by a prefix-free set of variable-length bitstrings. The assignment of symbols to bitstrings is determined by constructing a binary tree from the top-down, attempting to evenly partition the current set of symbols according to total frequency into two subsets at each node of the tree. Improvements on this greedy tree-based coding led to Huffman coding.

The state-of-the-art in compression speed remains Huffman coding (Collet, 2019), which is described in (Huffman, 1952) with the binary tree constructed in a bottom-up manner which takes into consideration the frequency of each symbol. This non-greedy method of tree-building was found to be an improvement over Shannon-Fano coding, resulting in a closer, but yet still imperfect, approximation to Shannon entropy. Since this construction operates at the symbol level, Huffman coding schemes cannot break a one-bit-per-symbol lower bound. Additionally, Huffman coding schemes require that a symbol be represented using an integer number of bits. Together, these limitations result in an imperfect approximation of the Shannon entropy, and therefore a limited compression ratio. While workarounds such as grouping multiple symbols together allows one to arbitrarily approach Shannon entropy, this results in a large alphabet size and thus a larger tree. These limitations apply in general to all symbol-level coding schemes, which can be considered as stateless (since there is no memory of previous symbols). Better compression ratios can only be obtained through the use of stateful or message-level coding schemes (Duda, 2013).

### 2.2 The dictionary problem in genetics

Dictionary compression methods are one approach towards stateful compression schemes. The *Zlib* (Gailly and Adler, 2004) and *Zstd* (Collet and Skibinski, 2019) methods used for the *bgen* or *ped* formats (Band and Marchini, 2018; Chang et al., 2015) are examples of dictionary methods. These methods generate the dictionary in an online fashion during the compression process, which decreases the compression rate. As the dictionary increases in size, queries into the dictionary become increasingly computationally intensive. While previously compiled dictionaries can be used in such methods to avoid online dictionary building, the compressed size of datasets which do not resemble those used to compile the dictionary will remain suboptimal, which is an issue given the diversity of genetic datasets. While dictionary methods provide a stateful compression scheme which operates above the symbol-level, these issues suggest the possibility of better performance from stateful and message-level coding schemes which do not use dictionaries. Indeed, *Zstd* itself uses *fse* to compress its dictionary, and the approach we adopt will simply omit the dictionary and use *fse* directly.

### 2.3 Arithmetic coding and the interval problem

Arithmetic coding methods are stateful, message-level, and dictionary-less coding schemes. Rather than assigning symbols to variable-length bitstrings, as in Huffman coding, or assigning sequences of symbols to bitstrings, as in dictionary methods, arithmetic coding methods assign entire messages to real numbers, with the goal of minimizing the length of the binary representation of the assigned number.

We briefly describe the high-level operation of a real-valued unit interval arithmetic encoder. Such an arithmetic coder maintains two state variables which represent a subinterval within the unit interval. Compression is achieved by a bijection between all possible messages and the real numbers within the unit interval, with messages of higher probability being assigned real numbers with shorter binary representations. Given an alphabet Σ, a symbol probability distribution 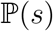 defined over all symbols *s* ∈ Σ, and a message *σ* = *σ*_1_ – *σ*_n_ ∈ Σ*, such a real number with short binary representation is found by contracting the current subinterval over *n* iterations such that the size of the *i* + 1 subinterval (as a proportion of the immediately preceding *i* subinterval) equals 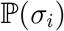. The shortest binary representation of a real number in the final subinterval is the encoding of *σ*. In practice, finite-precision floating-point arithmetic necessitates rescaling of the subinterval when possible, (*i.e*. doubling the interval when the result would remain a subset of the unit interval and outputting one bit of the binary representation).

The maintenance of two state variables to describe the current subinterval and the rescaling operations impose a computational cost to arithmetic coding. Such state maintenance operations could be reduced if one could reduce the number of state variables, and rescaling operations could be accelerated by replacing real intervals with integers.

### 2.4 Solutions to the interval problem

Asymmetric numerical systems provide a speed improvement over arithmetic coding by reducing two real state variables into one integer state variable and by replacing the representation of symbols using subintervals with one using uniformly distributed subsets which partition the integers. This replacement of real intervals with integer partitions and elimination of a state variable preserves the advantageous compression ratio of arithmetic coding while operating at a higher speed.

The additional speed improvement of *tans* methods (Duda, 2013) such as *fse* (Collet, 2019) arises from the implementation of the compression algorithm as a table (hence the “tabled” in *tans*) recording the transitions of a finite-state machine (hence the “finite state” in *fse*). This is a computationally efficient technique which has also found ubiquitous usage in fast string matching algorithms, such as modern regular expression engines. The speed of *fse* approaches Huffman coding, yet without the compression ratio issues. For these reasons, *fse* was chosen to be the compression method of choice for *agent* over all of the aforementioned techniques.

## 3 Methods

We conduct a series of experiments to explore the speed and size of the *finite state entropy* compression method on population genetics data through modifications of the open source *qctool* software package, and development of the *agent* software package.

### 3.1 Experiment 1: Partially exchangeable data

We examine the speed and compression ratio of our *agent* method in the partially exchangeable case. We consider data from the Thousand Genomes Project^1^. These data consist of 2,504 subjects sampled from 26 populations (sized between 61 and 113 samples) and 81,271,745 hard-called markers (The 1000 Genomes Project Consortium, 2015). We consider two presentations of these data. In the first presentation EXCH, subjects are ordered according to a uniformly random permutation and in the second presentation PART, subjects are ordered in a block structure such that the subjects in each population are ordered contiguously (*i.e*., all subjects from the first population appear first in the ordering, followed by all subjects from the second population, and so on). For each of these two presentations, we consider a task in which an uncompressed bgen file is recompressed using *fse* or *Zstd* compression algorithms. In each case, all of the Thousand Genomes Project data is presented in a single *bgen* file (*i.e*., with all chromosomes concatenated). So, for timings and compression sizes of *bgen* with *fse*, results are obtained by timing a version of *qctool* that has been modified to allow compression according to the *fse* algorithm.

### 3.2 Experiment 2: Larger studies

Considering the excellent performance of *plink2* in terms of speed and size displayed in Experiment 1 (enumerated in Table 1), we note that our *qctool* +*fse* patch outperforms the reference *qctool* +*Zstd* code. The reduced performance of *qctool +fse* with respect to *plink2* could be due to *qctool* overhead that is addressed by *plink2* and also small sample size considerations (*plink2* includes non-random access methods in their implementation of the LD-Compress algorithm Chang *et al*. 2019). Figure 1 indicates that *fse* may be indicated for sample sizes that are larger than the Thousand Genomes Project studied in Experiment 1 (and in the age of large-scale consortia, such sample sizes are of more interest). Motivated by these considerations, instead of implementing *fse* approaches to compression as a patch for *qctool*, we continue by developing a new software package implementing population-level variant-major compression and GWAS based on *fse*. We call this the *agent* software package, and this software package. We then conduct experiments with this *agent* software on larger sample sizes.

**Table 1:** Speed and size of Thousand Genome Project compression. Arithmetic coding of *bgen* files with *qctool* improves speed by 41% and size by 13% in the EXCH condition and improves speed by 45% and size by 16% in the PART condition. Note that *Zstd* compressed size is slightly larger in the PART condition, indicating that (counterintuitively) regularity in *bgen* files does not lead to improved compression ratios with *Zstd*. On the other hand, in this experiment compression size for *fse* does not depend on exchangeability.

| Method | EXCH speed(h) | PART speed(h) | EXCH size(Gb) | PART size(Gb) |
| --- | --- | --- | --- | --- |
| *qctool* + *Zstd* | 21.66 | 19.24 | 13.26 | 13.80 |
| *qctool* + *fse* | 12.83 | 10.54 | 11.55 | 11.55 |
| *plink2* | 0.79 | 0.78 | 5.92 | 5.90 |

In Experiment 2, we create biologically plausible simulated versions of the chromosome 22 data based on chromosome 22 from the Thousand Genomes Project with between 20,000 and 80,000 samples. This simulation is done using the *hapgen* software (Su et al., 2011). We create a single and independent simulated dataset for each number of samples (for each multiple of 10,000 between 20,000 and 80,000). After removing multi-allelic sites (a step required by *hapgen*), 1,055,452 variants remain on chromosome 22. We save each simulated dataset into a file encoded with the *bgen* format (using *Zstd*), and then consider a task in which *plink2* or *agent* is used to convert the *bgen* file into respectively the *ped* format (for *plink2*) or back to a version of the *bgen* format in which the *Zstd* compression algorithm for the compressed variant blocks of the *bgen* file are replaced by *fse* instead of *Zstd* (as in the *qctool+fse* condition of Experiment 1). The sizes of the resulting files and the runtime of the tasks are reported in Figure 3. Note that the number of variants is only a fraction of what is usually acquired in consortium style studies.

**Figure 2:**
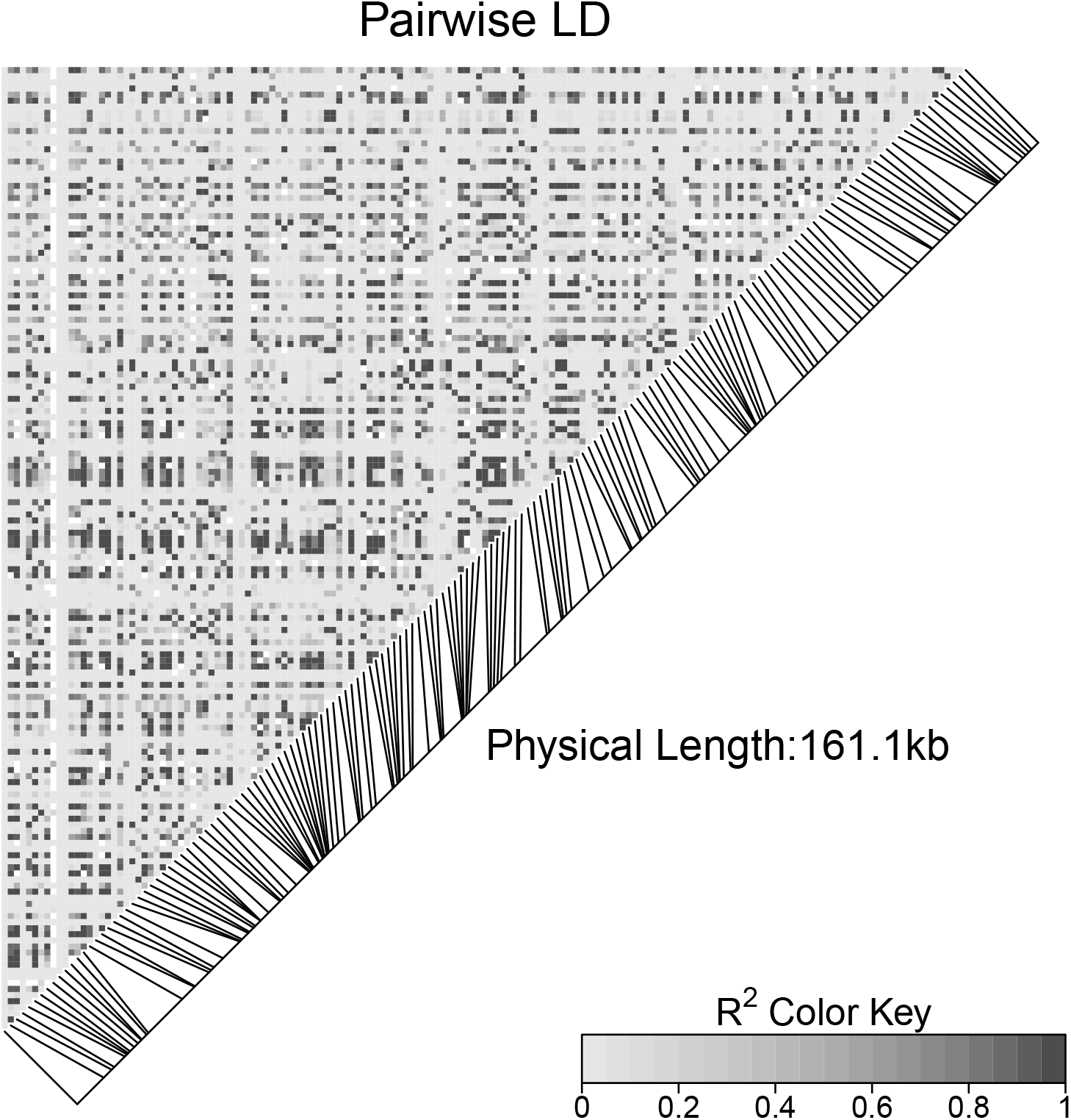
Linkage disequilibrium for a chunk in Experiment 3, displayed using *LDheatmap* (Shin et al., 2006).

**Figure 3:**
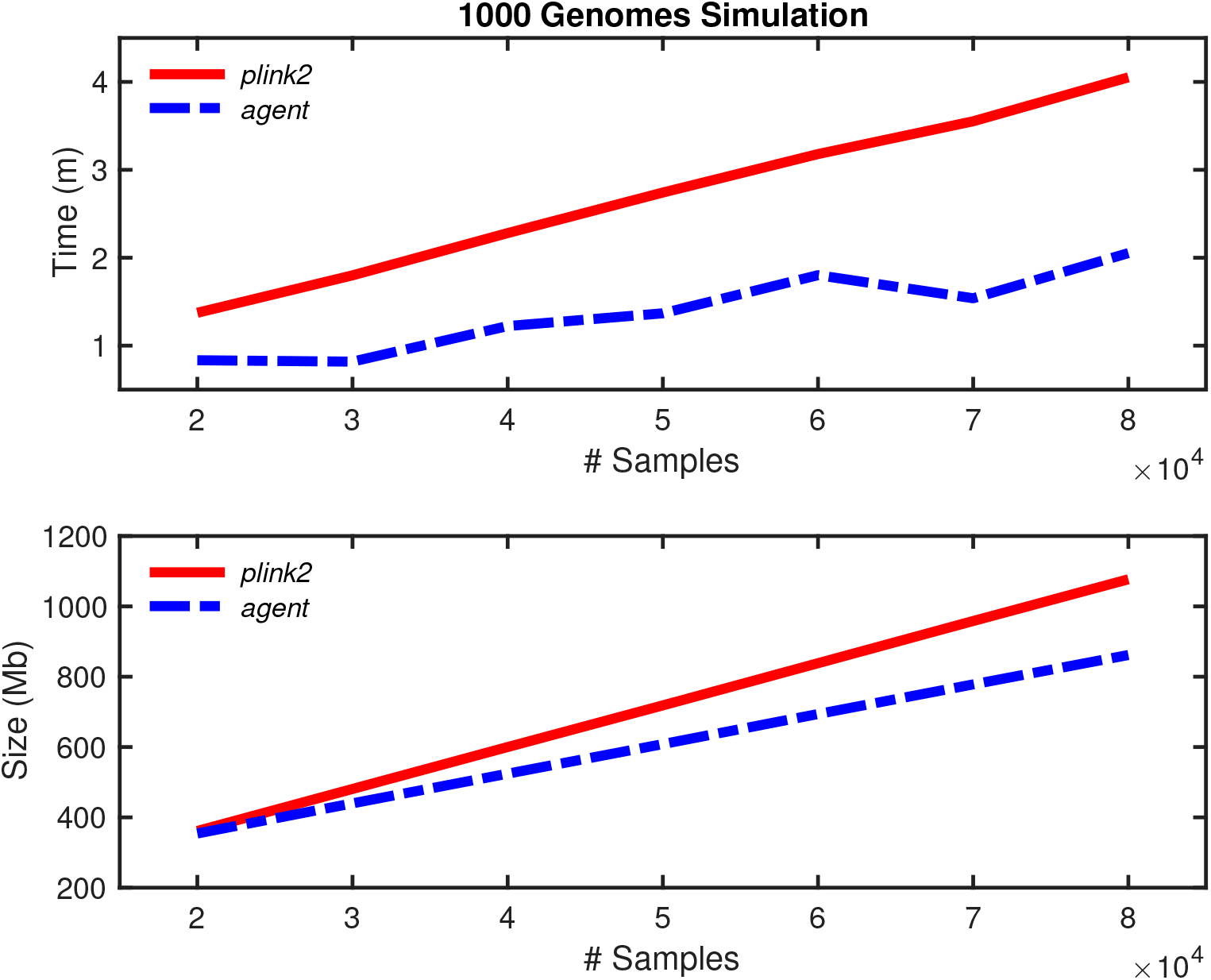
Performance of *plink2* versus *agent* in a task in which between 10,000 and 80,000 simulated samples are considered. Simulation is based on extending the Thousand Genomes Project to create a biologically plausible sample of chromosome 22 on many more subjects using the *hapgen* software. The *agent* method outperforms *plink2* for all conditions, but extrapolation of resulting file size indicates improved performance of *plink2* under small sample sizes (as indicated by Table 1 columns 4 and 5).

### 3.3 Experiment 3: Compressing dosages

Modern GWAS often rely on genotype imputation methods in which variants that are not typed in a study sample are imputed through use of dense reference panels (the first large GWAS considering this approach was Wellcome Trust Case Control Consortium 2007). Studies may also make use of genotype likelihood information in which the number of copies of an allele are expressed as a probability distribution that reflects uncertainty in the assay. In these two situations, genotypes are often expressed as a real number indicating the dosage: the expected number of minor alleles. While this results in loss of information about the phasing and the uncertainty in the genotype, *plink2* and other tools implementing univariate regression consider dosage, and in some cases the *ped* format stores dosages (discarding genotype uncertainty) in a fixed point 16-bit numerical representation (Chang *et al*., 2019).

We consider a simulation study in which dosages are compressed. We simulate 490,000 haplotypes under the coalescent with recombination with 100,000 segregating sites and 1,000 recombination sites over a number of basepairs roughly equivalent to chromosome 22 using the *ms* software (Hudson, 2002). We use 10,000 of the haplotypes as a reference panel, and we combine the remaining 240,000 haplotypes into diploid genotypes forming a study panel. For details on study and reference panels in imputation we refer to Howie et al. 2009. We vary the amount of imputation over the four conditions, and for each condition we remove X% of the markers for X% of the samples in the study panel, and then impute the removed genotypes using the reference panel. We vary *X* over the range 20%, 40%, 60% and 80%. The same set of markers is removed for each selected sample leading to the blockwise study/reference paradigm. We conduct the imputation using the *impute2* software. The impute2 software was run on 80 equally sized and partially overlapping chunks. A linkage disequilibrium plot is shown in Figure 2 for one of the chunks from this experiment.

In Figure 4, we consider a task in which chunks from the above described simulation are compressed with *Zstd* (using *plink2*) or with *fse* (using *agent*). The size of the compressed dataset and the speed of the compression are improved by *agent* under all imputation conditions. We note that compression using arithmetic codes is particularly recommended for dosage data stored in fixed point numerical representations with low bit rates. This is due to the highly skewed distributions over the genotypes that tend to result from imputation, as illustrated in Appendix A (these histograms are found from the default example provided with the *impute2* software). High skew allows arithmetic codes to efficiently assign codewords.

**Figure 4:**
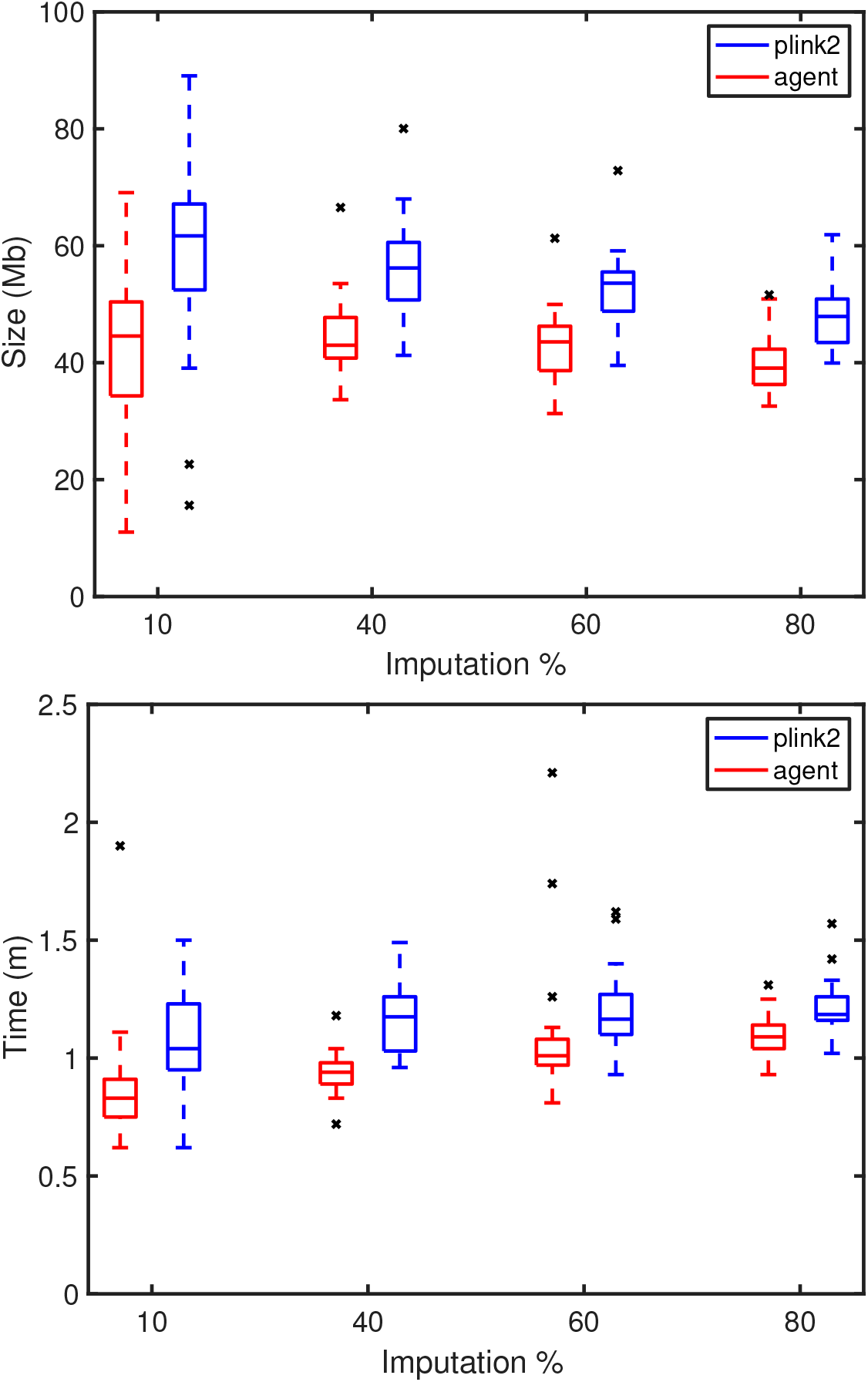
Compression speeds and sizes of genetic data including imputed dosages for 240,000 samples. Box plots indicate median and quantiles for 30 replicate datasets for each imputation condition. Datasets contain between X and Y (mean Z) variants.

### 3.4 Experiment 4: Genome-wide association study

To demonstrate the advantages of the *fse* methods displayed in the above experiments, we draft a request for comments on a new genetic file format supporting *fse* (the ‘.a1’ or ‘A one’ format), and provide the request for comments in Appendix B of the Supplementary Material. We implement two features for the *agent* software: 1) The conversion of *bgen* files to. *a1* files. 2) Univariate linear genome-wide association studies between. *a1* files (which may contain dosages) and multiple partially observed phenotypes. Then, we compare the *agent* software to a 2020 build of the state-of-the-art *plink2.00 software.*

We perform two examinations using 30/80 of the chunks for the simulations developed in Experiment 3. First, we consider 10 repeats for the 20% imputation condition and 512 phenotypes drawn independently from a standard Gaussian distribution (i.e.: *N* = 120, 000 samples, *D* = 512 phenotypes and *M* = 19, 922 of which 20% were imputed). We show box plots for the runtimes of the 10 repeats in Figure 5 (*Left*) and note that plink2 is only slightly faster than *agent* in this condition. In plink2, compression may not be done on dosages, which may explain the improved plink2 runtimes. Both pieces of software are optimized with Advanced Vector Extension (AVX2) instructions. Second, we consider the same genetic datasets and vary the number of phenotypes between 512 and 2048 (again, with independent standard Gaussian distributions). We perform one GWAS for each setting of the imputation proportion, yielding 5 runtimes for each method for each phenotype condition and these runtimes are plotted on Figure 5 (*Right*). This shows that the scalings of the runtime of *agent* and *plink2.00* as the number of phenotypes is varied are similar.

**Figure 5:**
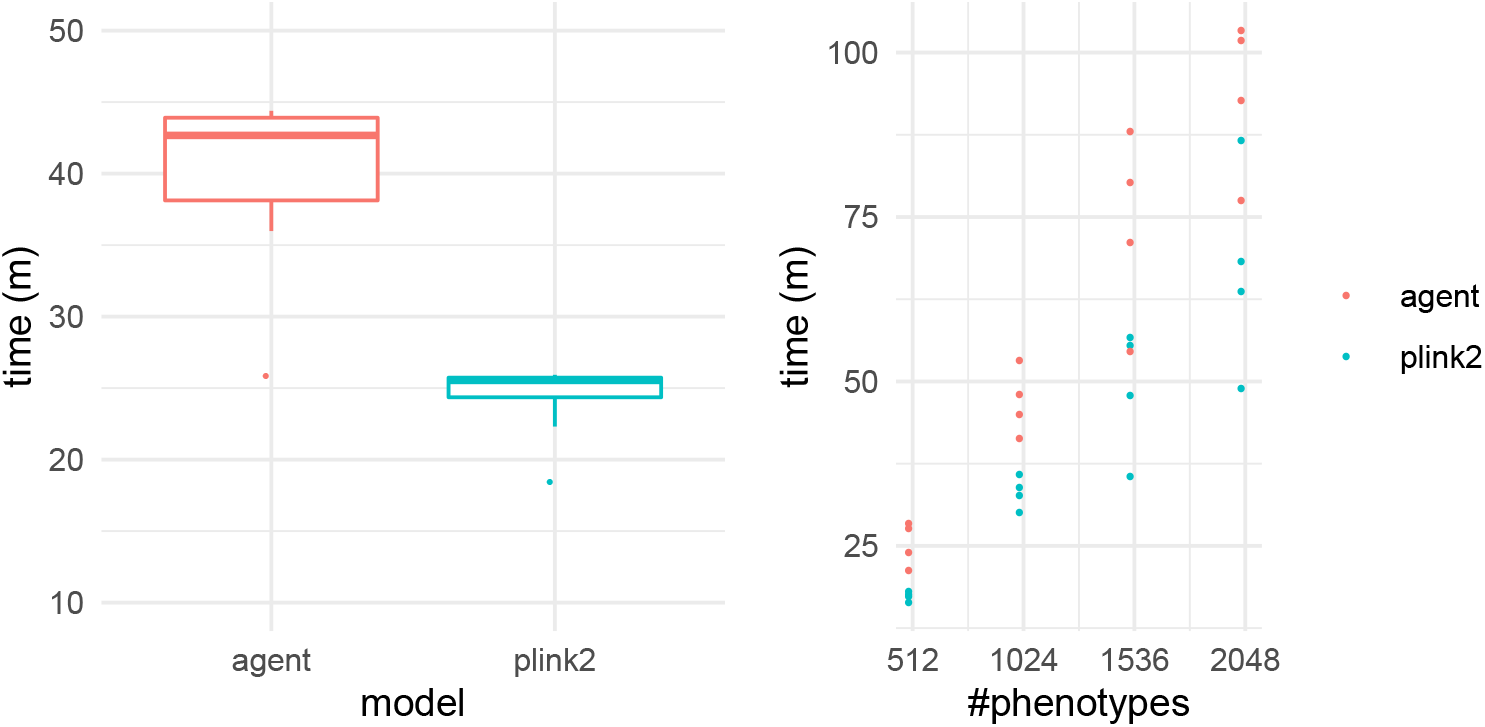
Speed comparisons between *plink2.00* and *agent* on imputed data. (*Left*) Ten independent restarts are considered for a dataset with 512 phenotypes. (*Right*) The number of phenotypes is varied. In each condition, *plink2.00* is only slightly faster than *agent*.

## 4 Discussion

We reviewed existing compression algorithms and their theoretical advantages and disadvantages for population genetic data storage, including the state-of-the-art *ans* and *tans* methods and *fse* compression. We noted the opposing compression ratio-speed tradeoffs in arithmetic coding versus Huffman coding and the ratio and speed issues with using existing dictionary methods such as *Zlib* and *Zstd* for population genetic data. We selected *fse* for its inheritance of the high compression ratio of arithmetic coding and the high compression speed of Huffman coding combined with the elimination of large and computationally costly dictionaries.

We compared compression time and compressed size for *fse* compression with existing methods of population genetic data storage over three experiments to test the effects of exchangeability, large number of samples and imputed dosages. In the fourth experiment we apply our *agent* tool using *fse* to a genome-wide association study and compare the time taken with the existing *plink2* tool.

In Experiment 1, a modified *qctool* with *fse* support compresses faster and smaller than *qctool*’s existing *Zstd* implementation for both fully and partially exchangeable data. In Experiment 2, our tool *agent* using *fse* compresses faster and smaller than the existing *plink2* tool on simulated chromosome 22 data with 1,055,452 variants over 20,000 to 80,000 samples (a typical dataset size in genetic analysis). In Experiment 3, *agent* compresses faster and smaller than *plink2* on imputed dosages for 240,000 samples over imputation percentages of 20%, 40%, 60% and 80%.

In Experiment 4, we add a GWAS feature to *agent* and show that its speed is only slightly slower than *plink2.00*. Since *fse* compression on population genetic data is faster than dictionary methods (as shown in Experiments 1 to 3 and Figure 1), the speed improvement in *plink2.00* is likely due to the high quality of the design of *plink2.00* (*i.e*., it’s likely that *plink2.00* speed would be improved should *fse* be adopted by *plink2.00* and also it’s likely that *agent*’s speed could be improved through improvements to the design). We also note that the relative speed between *agent* and *plink2.00* may differ for situations with more SNPs (we only examined a simulated region around 3/8ths of the size of chromosome 22, and the genotypes can still be stored uncompressed in Random Access Memory (RAM), which may positively influence the speed of *plink2.00*).

## 5 Conclusion

Through our review of entropy coding and dictionary compression methods and their relevant advantages and disadvantages for population genetic data, we settle upon *fse* compression, part of the larger *tans* and *ans* families of compression algorithms, for use in a novel population genetic data tool and format: *agent*. This tool compresses faster and provides smaller file sizes than existing methods for population genetic data. We demonstrate superior performance in both time and space domains for *fse* compression when compared to the existing methods of population genetic data storage regardless of exchangeability, large number of samples or imputed dosages over three experiments.

We apply *agent* to a genome-wide association study in a fourth experiment. With the recent rise of genome-wide association studies and the resulting needs for computational resources in both time and space domains, the improvements brought by *agent* will increase the speed and feasibility of population genome-wide association studies and reduce the cost of such studies for both public and private sector research. Ultimately, *agent* will assist in the speedy delivery of the results of such studies, which have real-world outcomes in the form of new medical treatments, diagnostics, and a better understanding of the human genetic code.

## Acknowledgements

We thank Alex Sweeten for helpful discussion, and we thank Fred Popowich and the Big Data Hub at Simon Fraser University for help with computational resources.

## Footnotes

1 The Thousand Genomes Project Phase 3: May 27th 2015 release.

