## Supplementary Material for "Compression for population genetic data through finite-state entropy"

### Efficient compression and GWAS for population genetic data

### Appendix A: Histograms of dosages

In this Appendix, we provide histograms for dosages computed using the *impute2* software (Howie, Donnelly, & Marchini, 2009). We consider the example genetic dataset provided along with the *impute2* software download and impute the missing genotypes in that dataset using the default *impute2* parameters. For each genetic marker, we provide a two-dimensional histogram. The scales of the dimensions correspond to the proportion of the reference and alternate allele (respectively). All of the markers are bivariate. These histograms indicate that imputed genetic data are sparse.

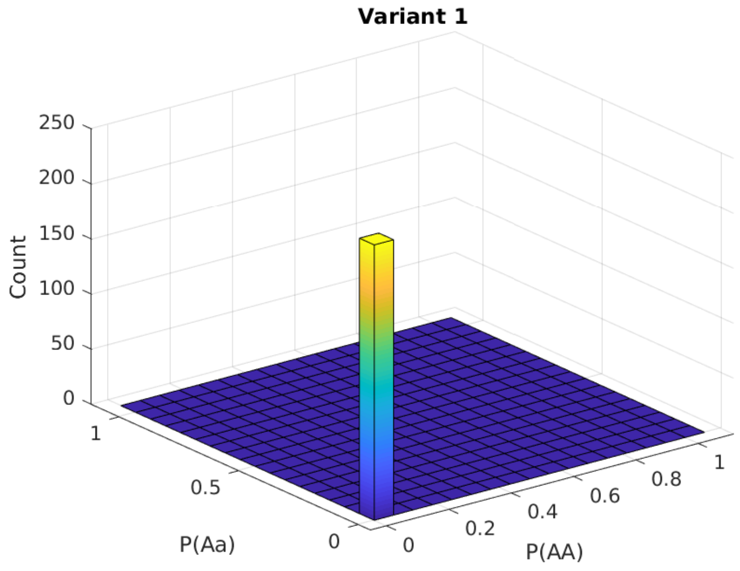

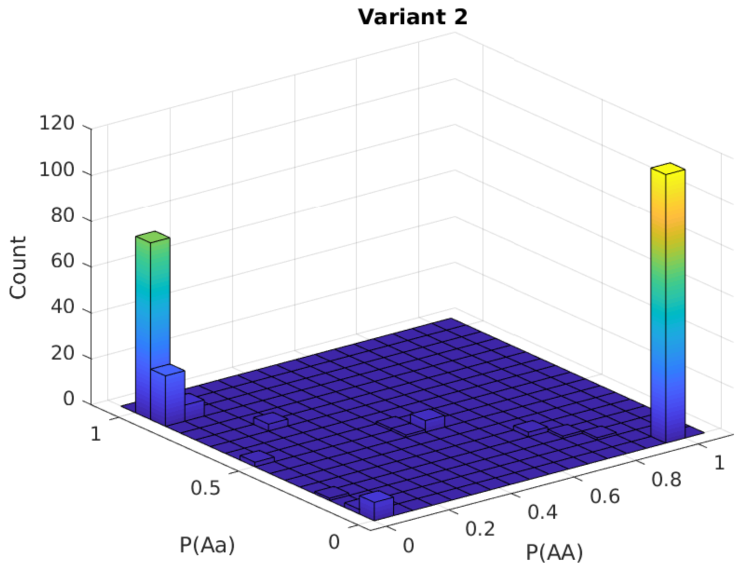
Variant 2
Count
120
100
80
60
40
20
0
1
0.5
0
P(Aa)
0
0.2
0.4
0.6
0.8
1
P(AA)

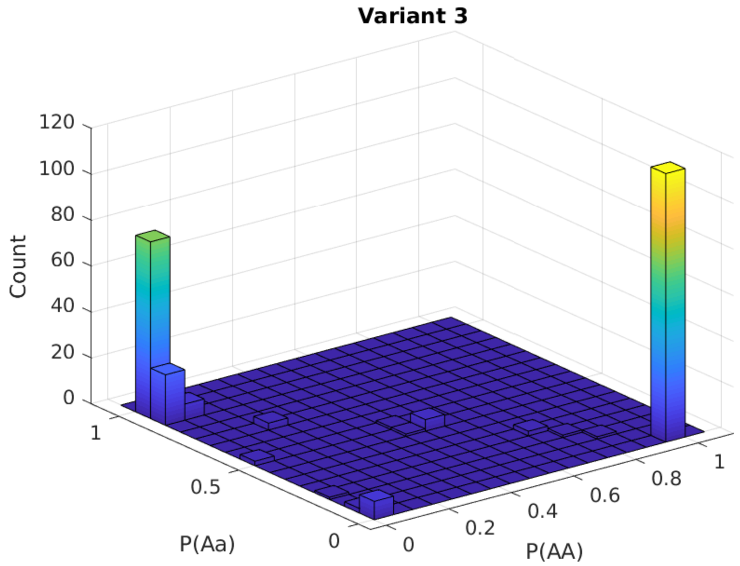
Variant 3
Count
120
100
80
60
40
20
0
1
0.5
0
P(Aa)
0
0.2
0.4
0.6
0.8
1
P(AA)

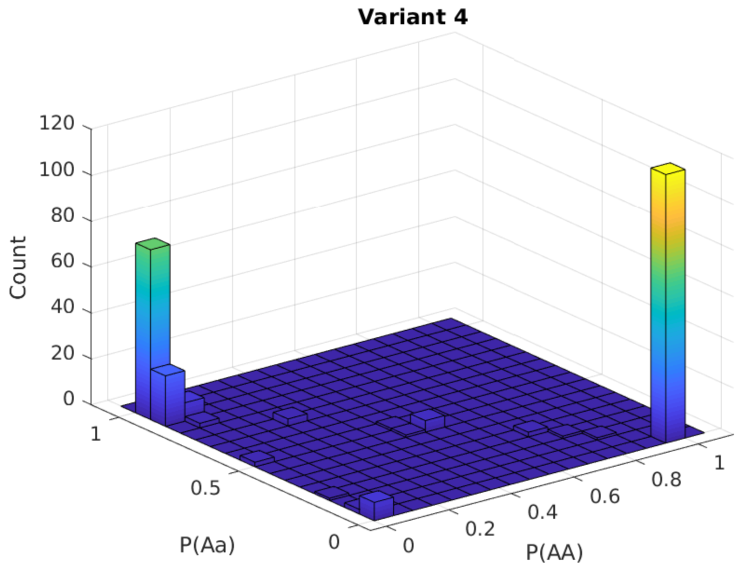
Variant 4
Count
120
100
80
60
40
20
0
1
0.5
0
P(Aa)
0
0.2
0.4
0.6
0.8
1
P(AA)

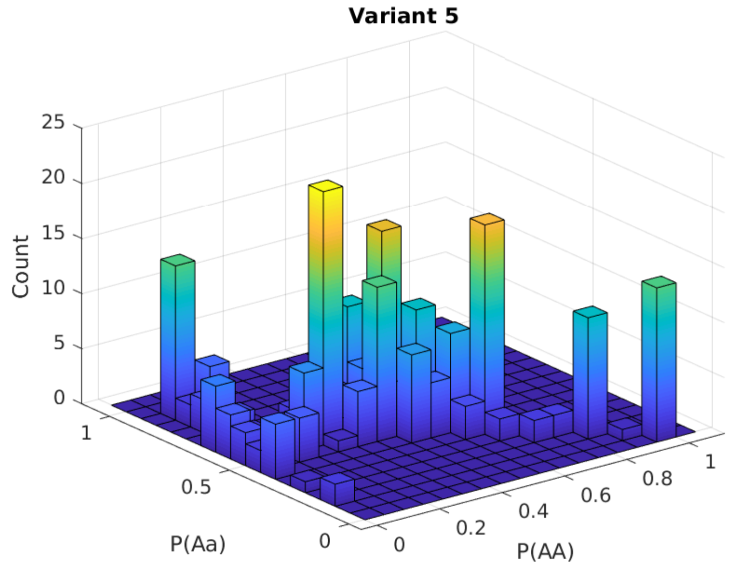
Variant 5
Count
25
20
15
10
5
0
1
0.5
0
P(Aa)
0
0.2
0.4
0.6
0.8
1
P(AA)

Variant 6

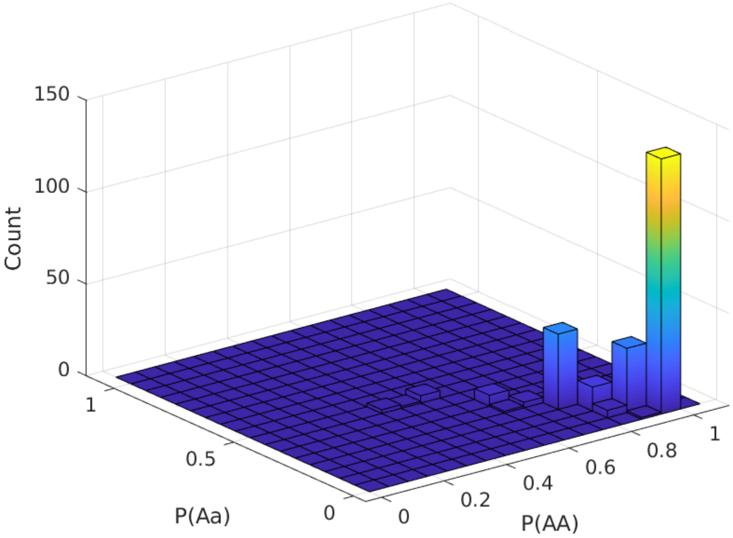


Variant 7

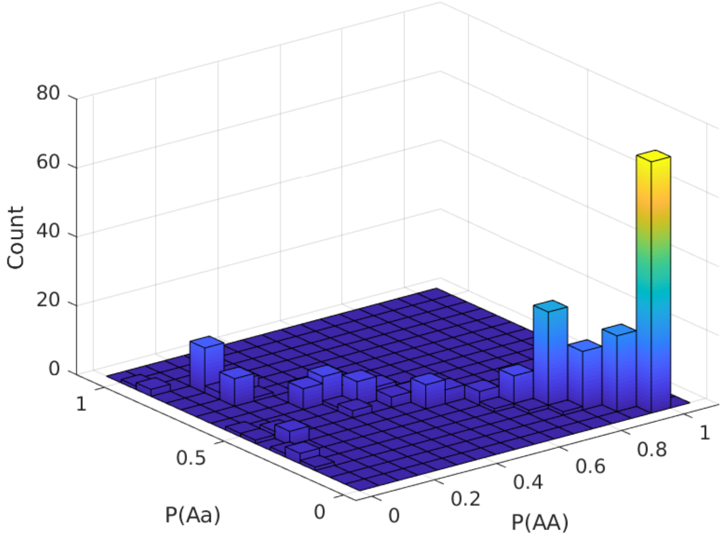

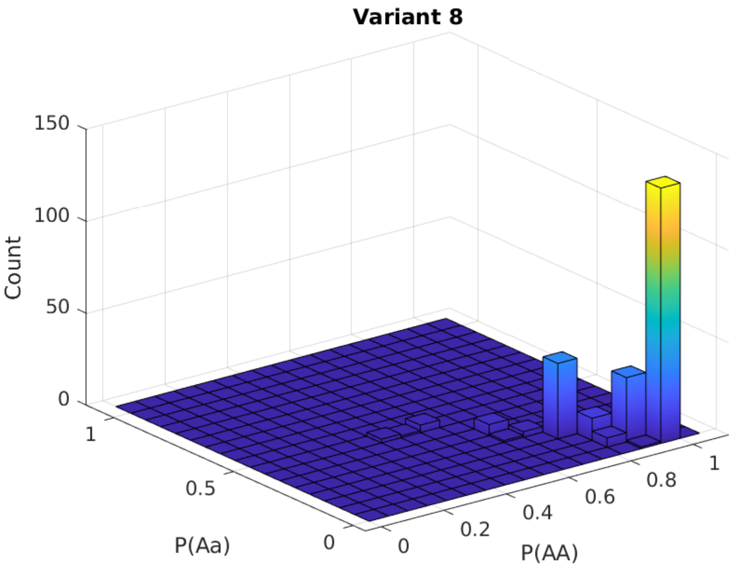
Variant 8
150
100
50
0
Count
1
0.5
0
P(Aa)
0
0.2
0.4
0.6
0.8
1
P(AA)

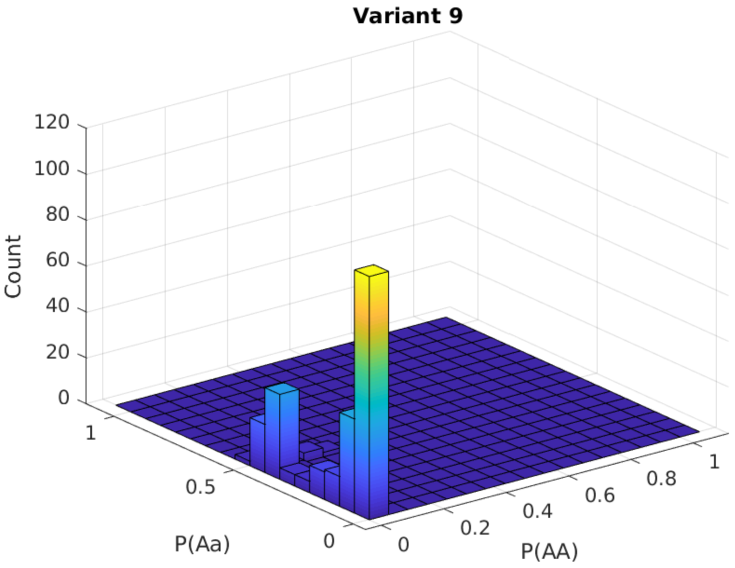
Variant 9
120
100
80
60
40
20
0
Count
1
0.5
0
P(Aa)
0
0.2
0.4
0.6
0.8
1
P(AA)

**Variant 10**

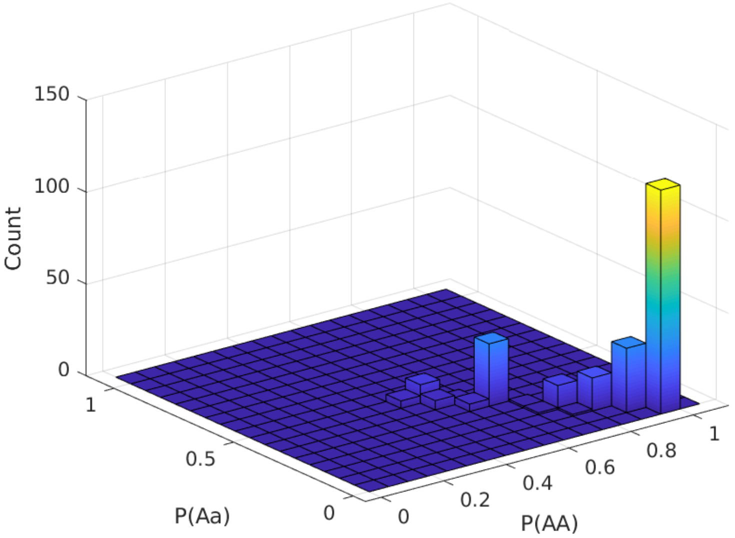


**Variant 11**

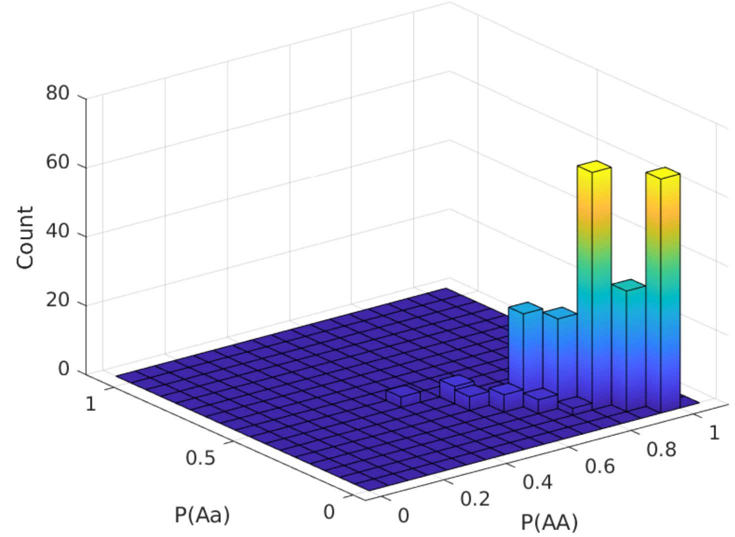

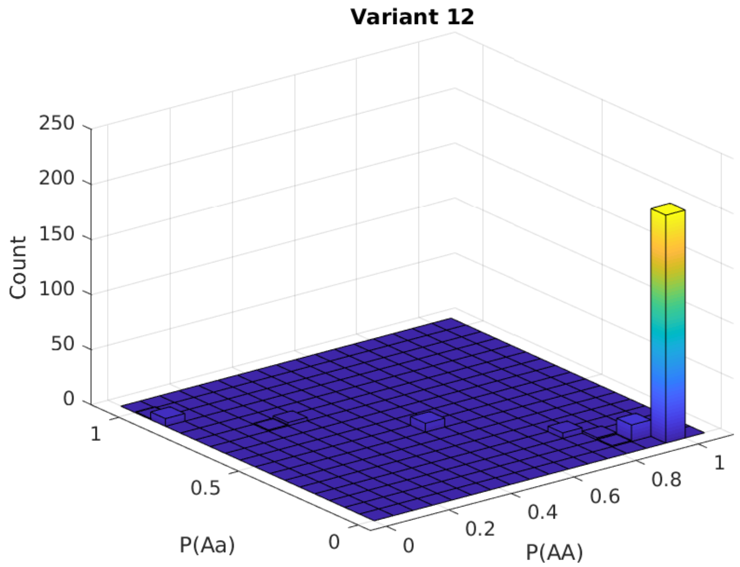
Variant 12
Count
250
200
150
100
50
0
P(Aa)
1
0.5
0
P(AA)
0
0.2
0.4
0.6
0.8
1

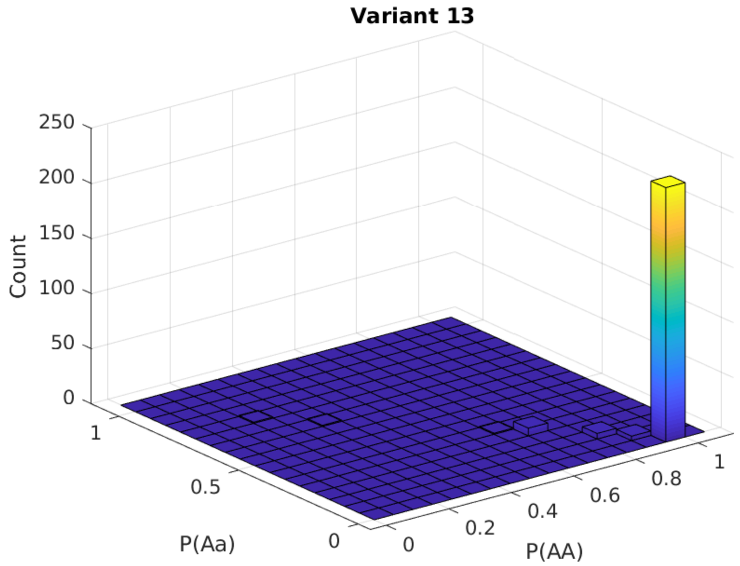
Variant 13
Count
250
200
150
100
50
0
P(Aa)
1
0.5
0
P(AA)
0
0.2
0.4
0.6
0.8
1

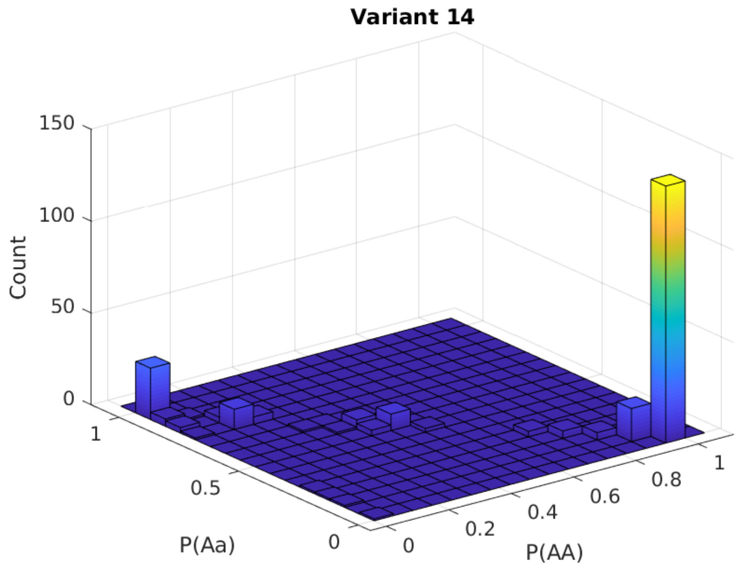
Variant 14
Count
150
100
50
0
1
0.5
0
P(Aa)
0
0.2
0.4
0.6
0.8
1
P(AA)

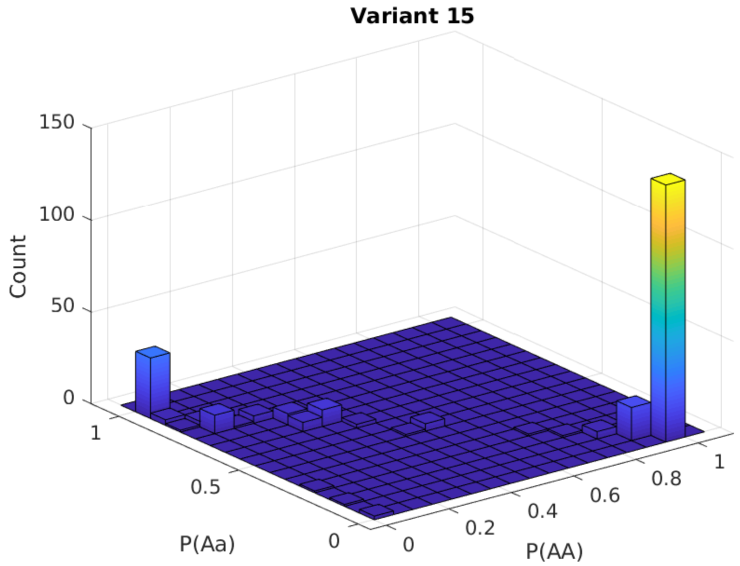
Variant 15
Count
150
100
50
0
1
0.5
0
P(Aa)
0
0.2
0.4
0.6
0.8
1
P(AA)

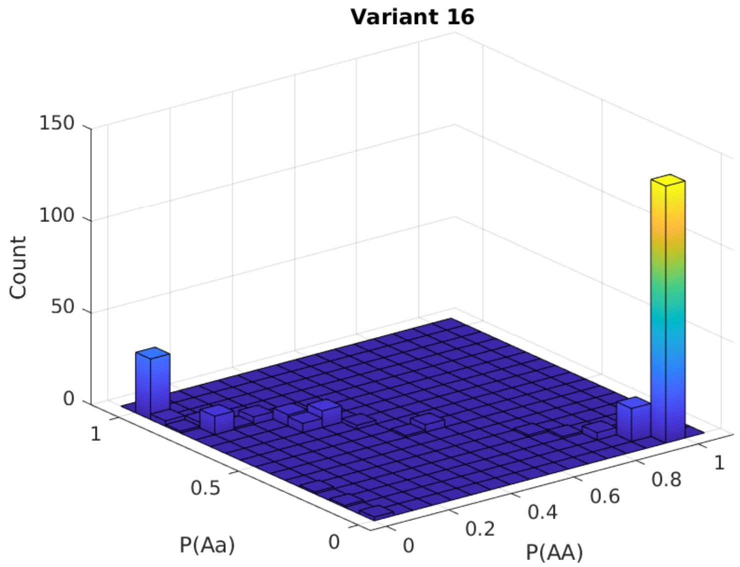
Variant 16
150
100
50
0
Count
1
0.5
0
P(Aa)
0
0.2
0.4
0.6
0.8
1
P(AA)

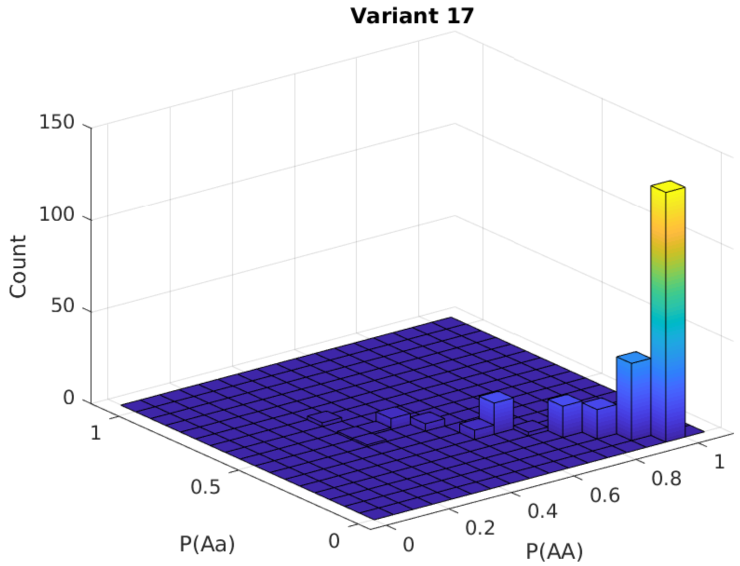
Variant 17
150
100
50
0
Count
1
0.5
0
P(Aa)
0
0.2
0.4
0.6
0.8
1
P(AA)

**Variant 18**

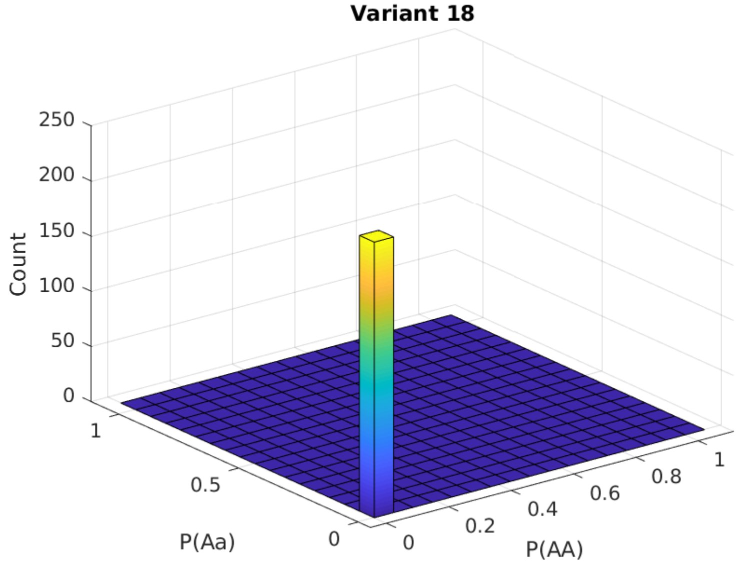


**Variant 19**

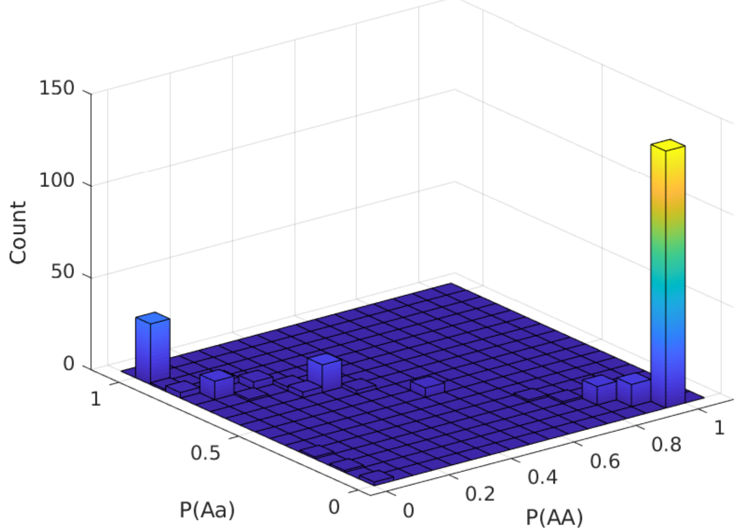

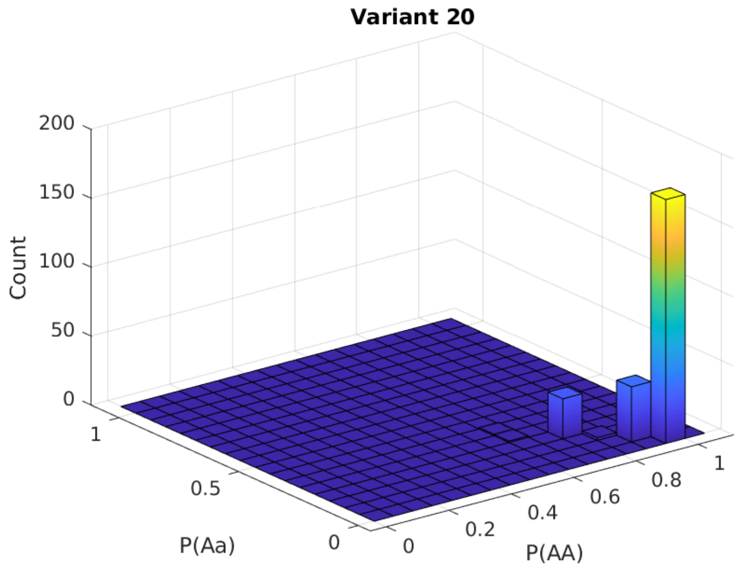
Variant 20
200
150
100
50
0
Count
1
0.5
0
P(Aa)
0
0.2
0.4
0.6
0.8
1
P(AA)

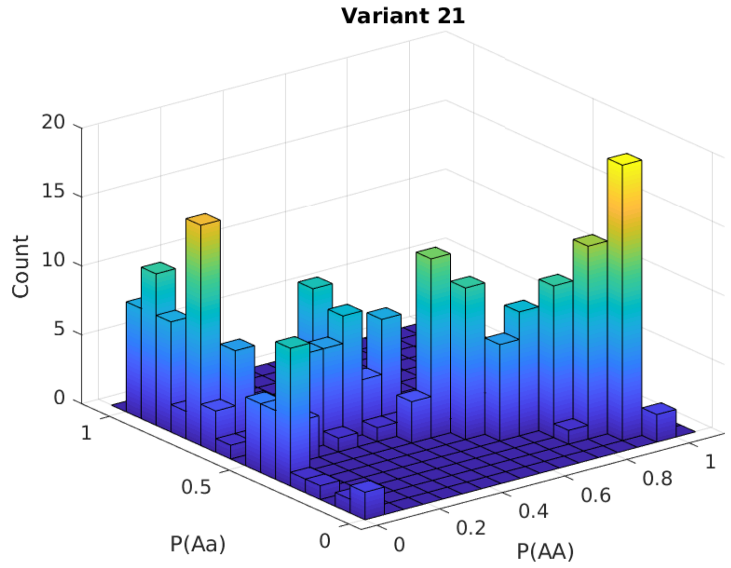
Variant 21
20
15
10
5
0
Count
1
0.5
0
P(Aa)
0
0.2
0.4
0.6
0.8
1
P(AA)

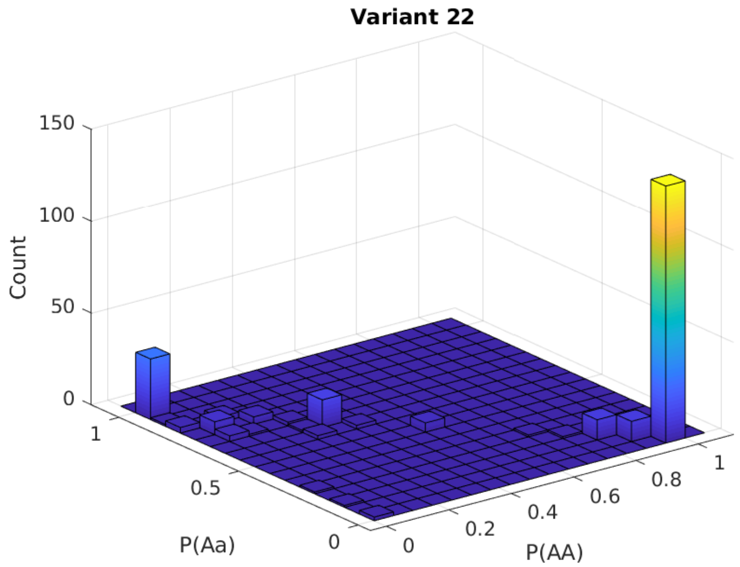

Variant 22
150
100
50
0
Count
1
0.5
0
P(Aa)
0
0.2
0.4
0.6
0.8
1
P(AA)


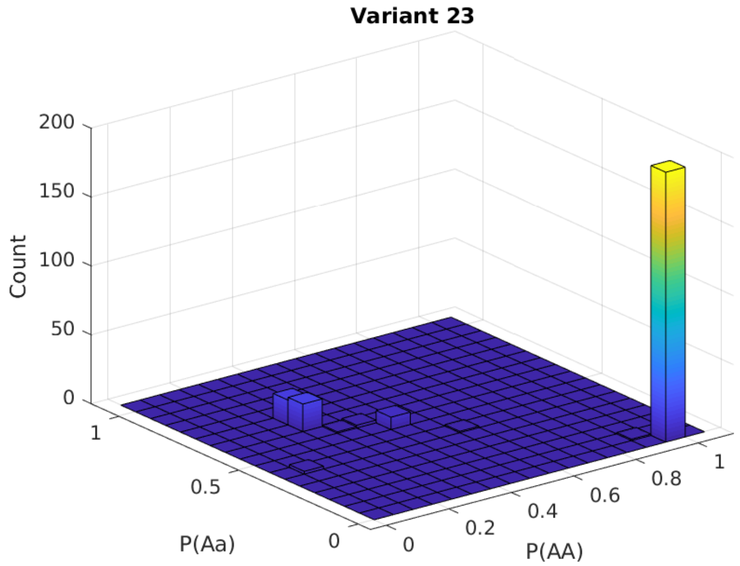

Variant 23
200
150
100
50
0
Count
1
0.5
0
P(Aa)
0
0.2
0.4
0.6
0.8
1
P(AA)

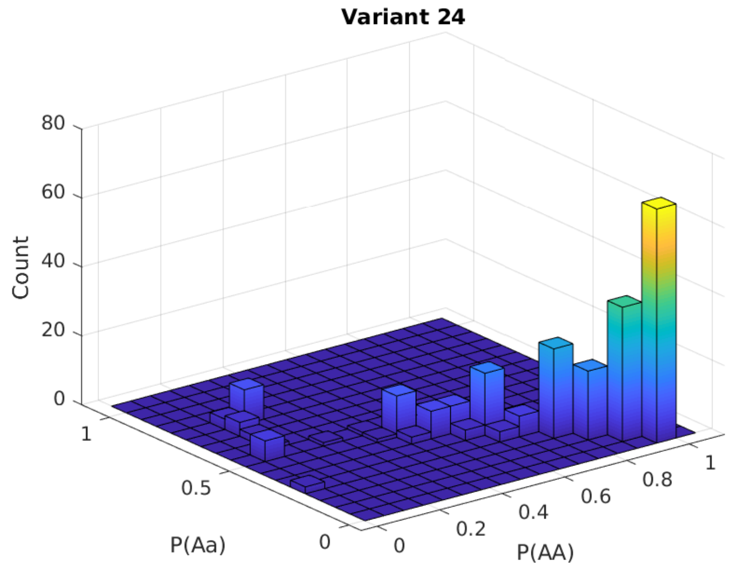
Variant 24
80
60
40
20
0
Count
1
0.5
0
P(Aa)
0
0.2
0.4
0.6
0.8
1
P(AA)

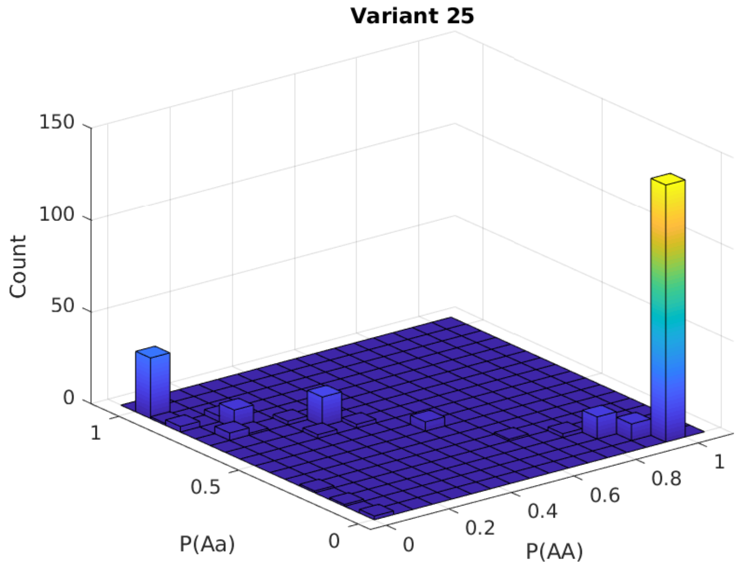
Variant 25
150
100
50
0
Count
1
0.5
0
P(Aa)
0
0.2
0.4
0.6
0.8
1
P(AA)

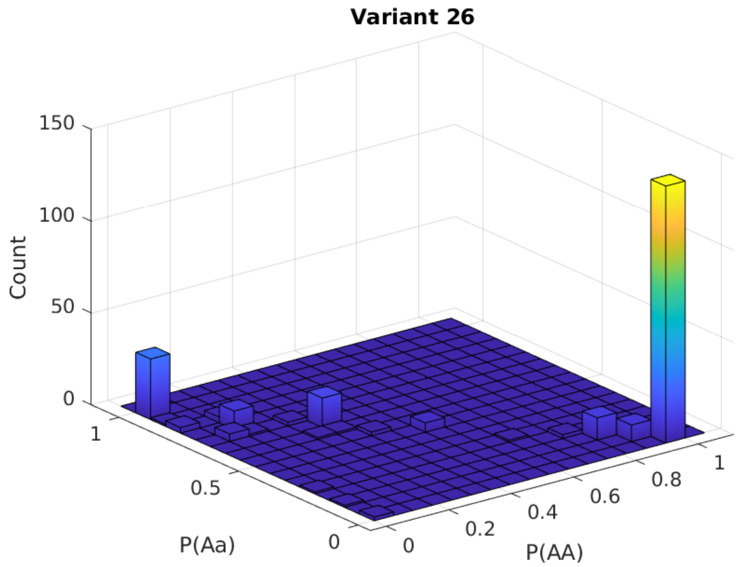
Variant 26
150
100
50
0
Count
1
0.5
0
P(Aa)
0
0.2
0.4
0.6
0.8
1
P(AA)

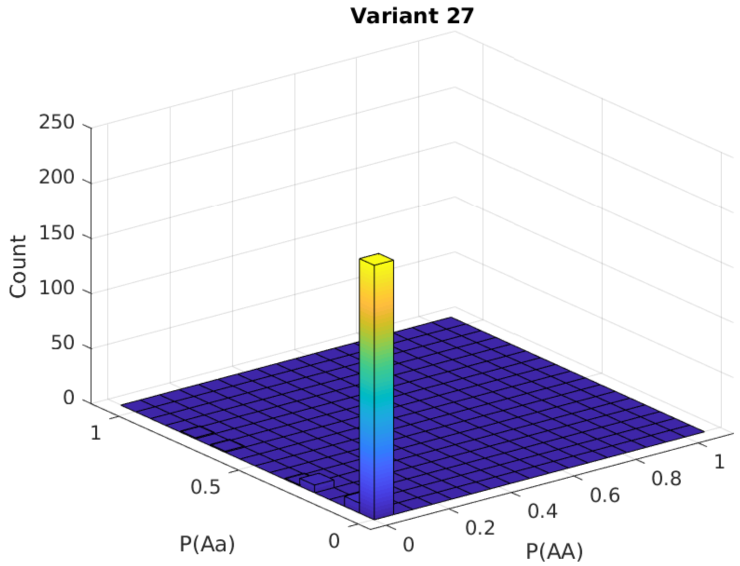
Variant 27
250
200
150
100
50
0
Count
1
0.5
0
P(Aa)
0
0.2
0.4
0.6
0.8
1
P(AA)

**Variant 28**

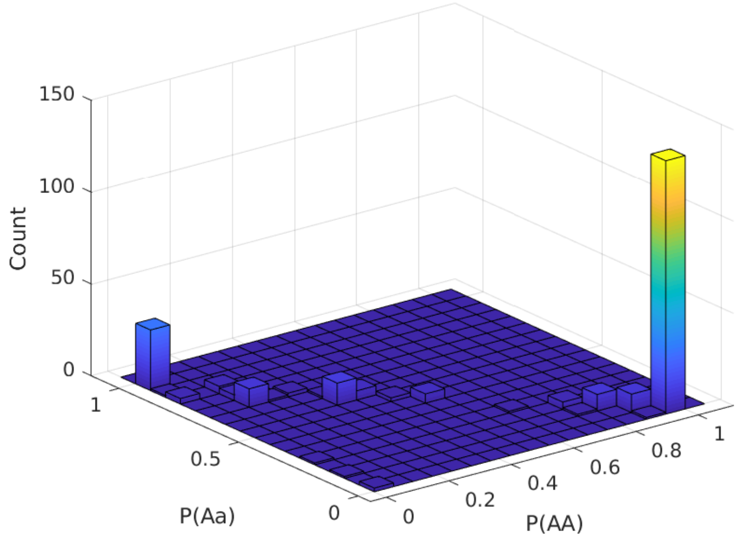


**Variant 29**

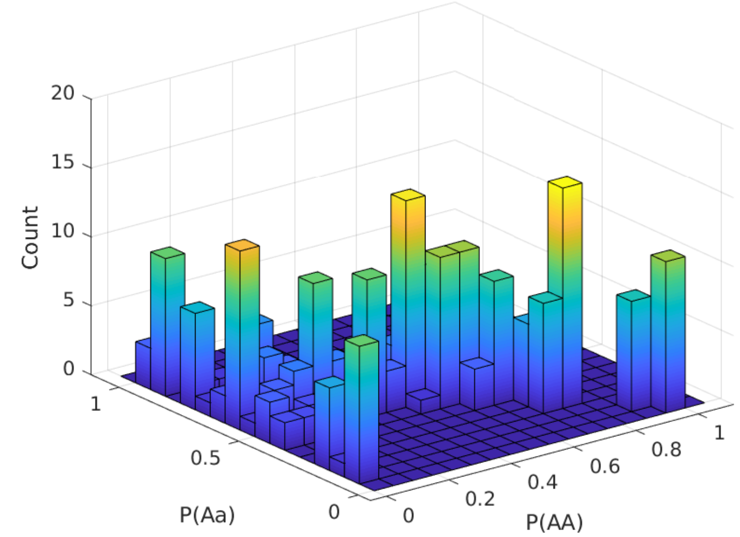

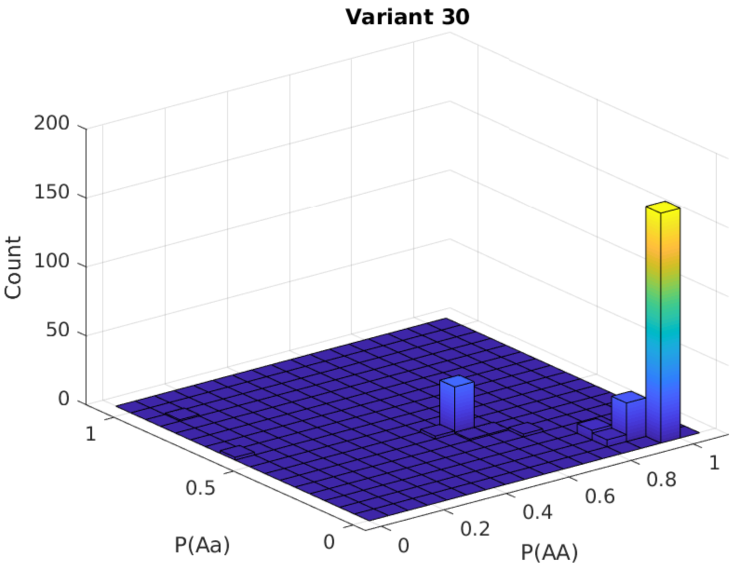
Variant 30
200
150
100
50
0
Count
1
0.5
0
P(Aa)
0
0.2
0.4
0.6
0.8
1
P(AA)

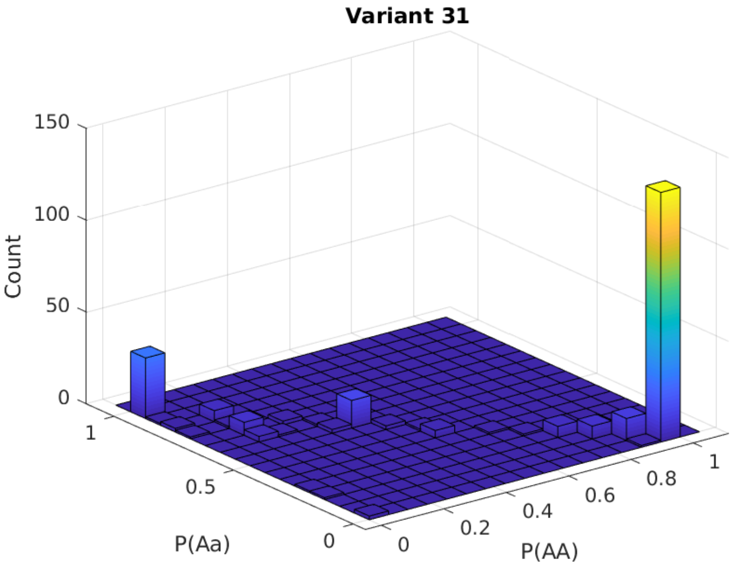
Variant 31
150
100
50
0
Count
1
0.5
0
P(Aa)
0
0.2
0.4
0.6
0.8
1
P(AA)

##### Variant 32

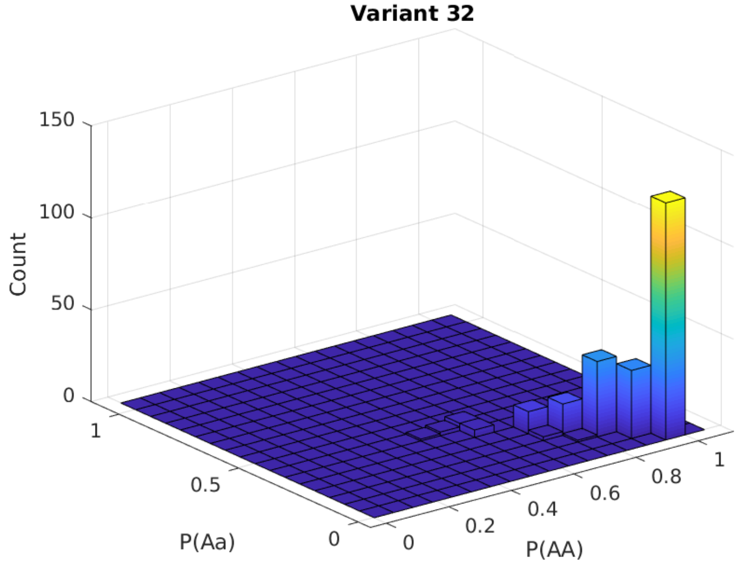


##### Variant 33

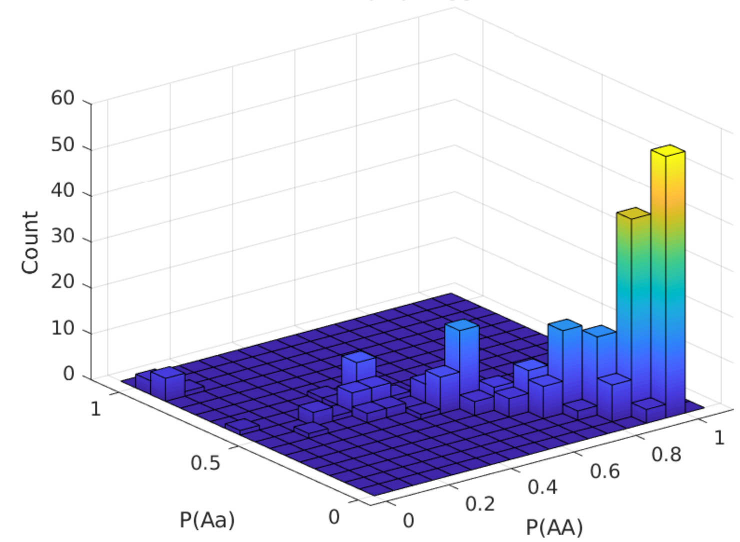

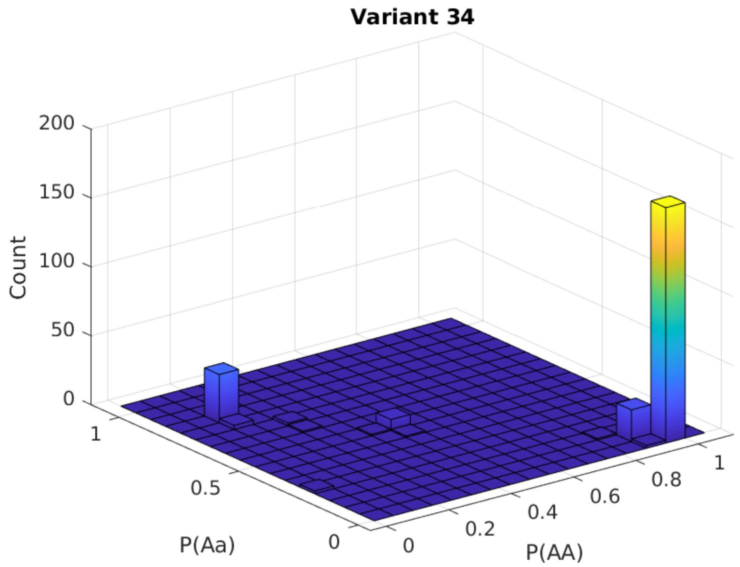
Variant 34
Count
200
150
100
50
0
1
0.5
0
P(Aa)
0
0.2
0.4
0.6
0.8
1
P(AA)

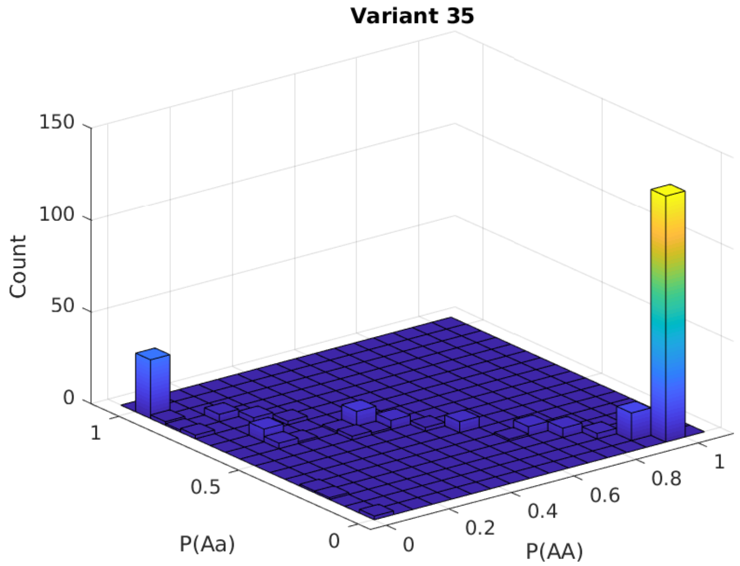
Variant 35
Count
150
100
50
0
1
0.5
0
P(Aa)
0
0.2
0.4
0.6
0.8
1
P(AA)

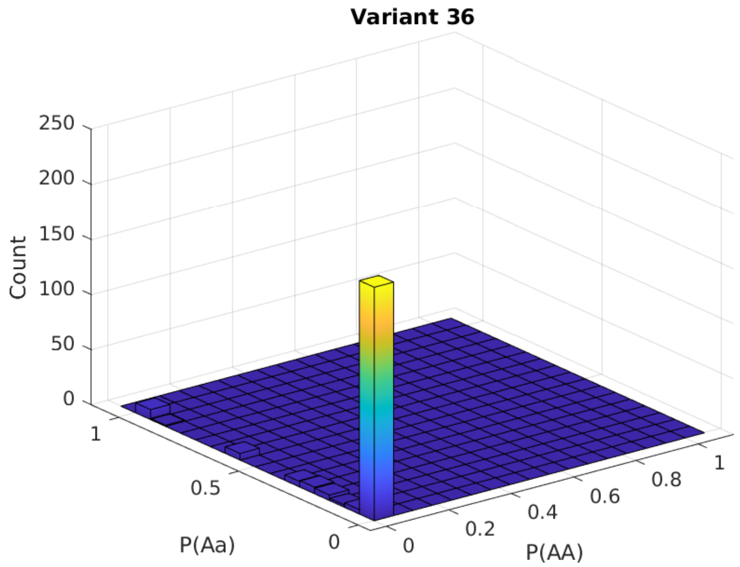
Variant 36
250
200
150
100
50
0
Count
1
0.5
0
P(Aa)
0
0.2
0.4
0.6
0.8
1
P(AA)

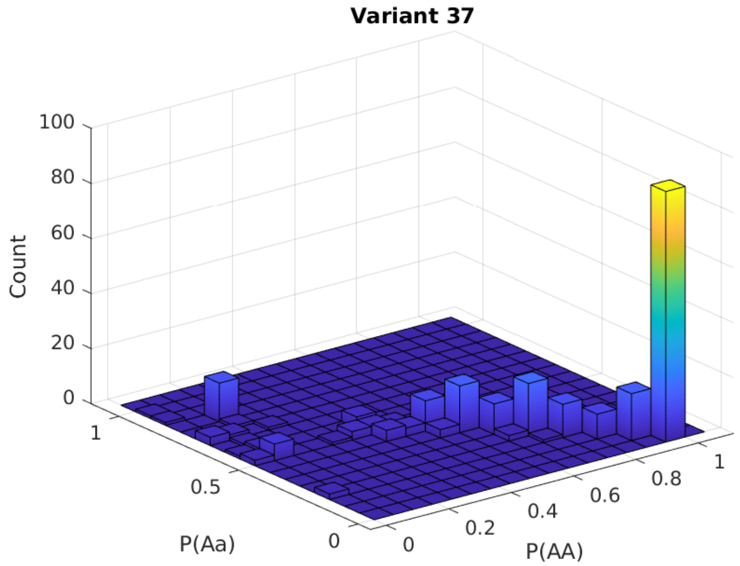
Variant 37
100
80
60
40
20
0
Count
1
0.5
0
P(Aa)
0
0.2
0.4
0.6
0.8
1
P(AA)

**Variant 38**

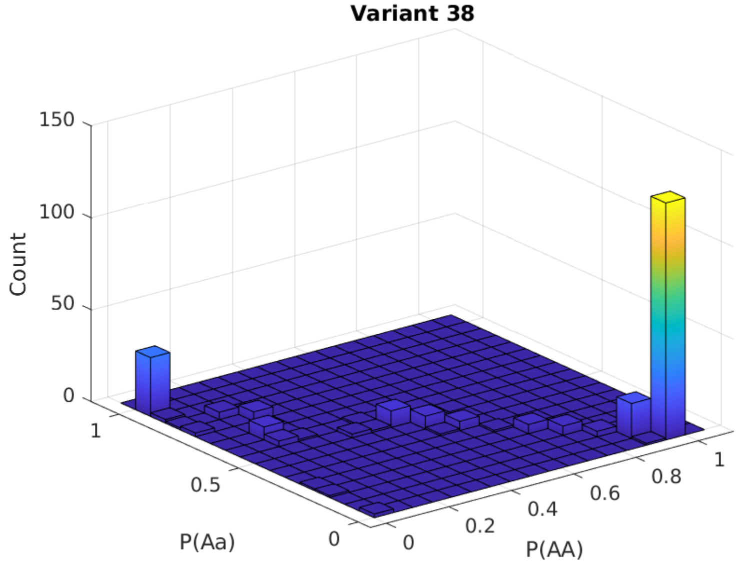


**Variant 39**

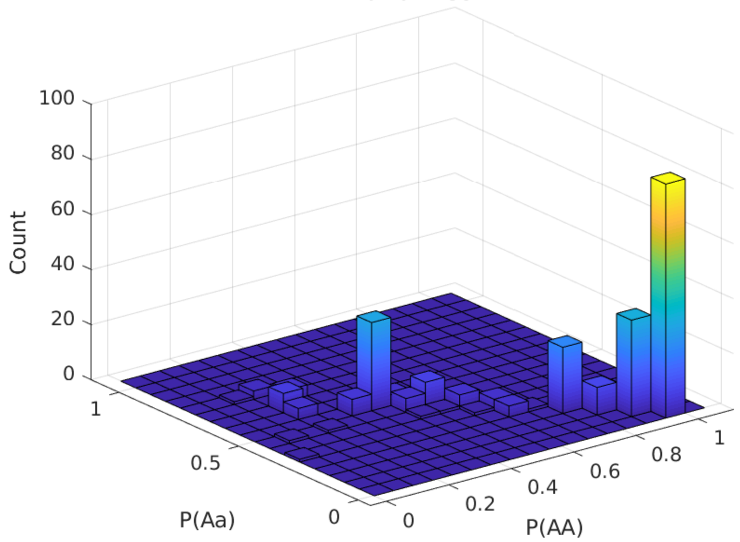

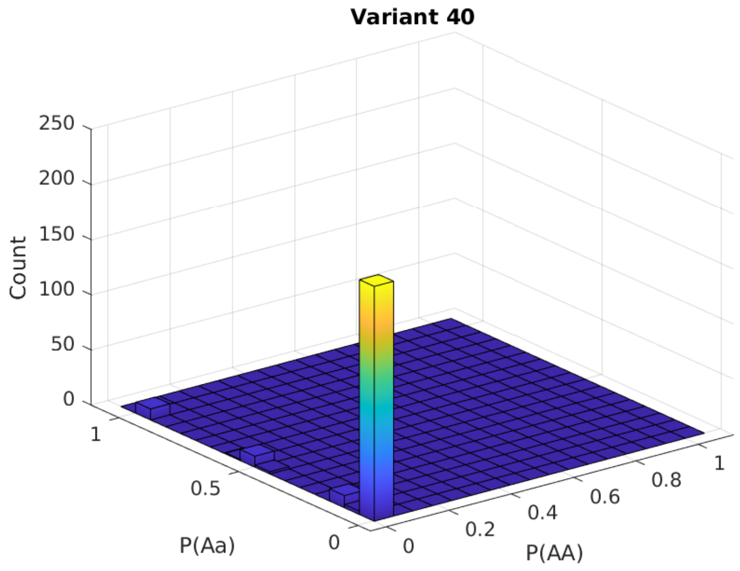
Variant 40
250
200
150
100
50
0
Count
1
0.5
0
P(Aa)
0
0.2
0.4
0.6
0.8
1
P(AA)

### Appendix B

##### *AGENT FORMAT REQUEST FOR COMMENTS*

Files encoded in this format must have a filename that ends in `.a1` (example: `chr01.a1`). All numbers encoded in `.a1` file must be in little-endian IEEE 64-bit formats.

**Goal**: Provide a minimal file format for demonstrating FSE compression, allowing fast GWAS.

- Header: (Length: *64 + 8 * M bytes*).
  - First 8 bytes. *(0)*. The ASCII strings `agent` (magic) and then `001` (version).
  - Next 8 bytes. *(8)*. The number of samples *N*, `uint64`.
  - Next 8 bytes. *(16)*. The number of variants *M*, `uint64`.
  - Next 32 bytes. *(24)*. Currently unused.
  - Next 8 bytes. *(56)*. *L0*: The absolute offset to beginning of variant block *1*. This is the number *64 + M * 8*, `uint64`.
  - For *i = 1 … M*:
    - Next 8 bytes. *(64 + 8 * i)*. *Li*: The absolute offset to end of variant block *i*, `uint64`.

- Blocks: (Length: *LM – L0*).
  - For *i = 1 … M*:
    - Next *X* bytes. *(L(i-1))*. Variant block. *N* dosages FSE compressed. The range of this block is given by *L(i-1)* and *Li* (left inclusive, right exclusive).

- Footer: (Length *Z*).
  - Next *Z* bytes. *(LM)*. Currently unused.
